# Niacinamide enhances cathelicidin mediated SARS-CoV-2 membrane disruption

**DOI:** 10.1101/2022.08.26.505399

**Authors:** Tanay Bhatt, Sneha Uday Khedkar, Binita Dam, Sahil Lall, Subhashini Pandey, Sunny Kataria, Paul M Dias, Morris Waskar, Janhavi Raut, Varadharajan Sundaramurthy, Praveen Kumar Vemula, Naresh Ghatlia, Amitabha Majumdar, Colin Jamora

## Abstract

The continual emergence of new SARS-CoV-2 variants threatens the effectiveness of worldwide vaccination programs and highlights the need for complementary strategies for a sustainable containment plan. A promising approach is to mobilize the body⍰s own antimicrobial peptides (AMPs), to combat SARS-CoV-2 infection and propagation. We have found that human cathelicidin (LL37), an AMP found at epithelial barriers as well as in various bodily fluids, has the capacity to neutralise multiple strains of SARS-CoV-2. Biophysical and computational studies indicate that LL37⍰s mechanism of action is through the disruption of the viral membrane. This antiviral activity of LL37 is enhanced by the hydrotropic action of niacinamide, which may increase the bioavailability of the AMP. Interestingly, we observed inverse correlation between LL37 levels and disease severity of COVID-19 positive patients, suggesting enhancement of AMP response would be an effective therapeutic avenue to mitigate disease severity and overcome vaccine escape.

---

SARS-CoV-2 infects host cells by binding of its spike protein to the Ace2 receptor on the cell membrane^1^. The importance of this interaction underlies the strategy of many vaccines to target Spike protein and thus prevent infection^2^. While exogenous Ace2 expression is sufficient to render cells competent for SARS-Cov-2 infection^3^, tissue expression of Ace2 is not always an indicator of viral tropism^4^. A case in point is the skin that expresses Ace2 in the epidermis in-vivo^5^, and keratinocytes in-vitro (Fig S1A) but nevertheless is not considered as a primary route of infection^6^. In support of this, exposure of human epidermal keratinocytes to SARS-CoV-2 does not result in a productive infection (Fig S1B). Thus, although epidermal keratinocytes are competent for SARS-CoV-2 infection, they may possess an endogenous defence mechanism to inhibit viral infection. The skin possesses a basal defence mechanism endowed by the constitutive secretion of antimicrobial peptides (AMPs)^7^. In particular, the human AMP cathelicidin (LL37) has been shown to target various classes of viruses^8,9^, including respiratory viruses^10^. Skin keratinocytes exhibit a higher basal level of LL37 secretion than lung epithelial cells (Fig S1C), which could be one of the factors leading to their lower infectivity by SARS-CoV-2 (Fig S1B).

To ascertain whether LL37 is effective against SARS-CoV-2, we incubated the virus with this AMP and assessed its ability to infect an intestinal epithelial cell (Caco2) as a reporter. We observed a dose-dependent decrease in viral gene expression upon treatment with LL37 (Fig 1A). This LL37 mediated effect was also observed in other SARS-CoV-2 variants (Alpha, Kappa, Delta, and Omicron) (Fig 1B). These observations were confirmed by tissue-culture infectious dose (TCID50) assays (Fig S1D). Previous work has suggested that LL37 can interact with the spike protein and the Ace2 receptor and possibly occlude the interaction surface between them^11^. However, LL37 has also been shown to interact with, and aggregate on membranes^12^. Hence, we hypothesized that LL37 may inhibit viral infection in a Spike/Ace2 independent manner. We thus compared the neutralising capacity of LL37 against viruses of different tropism, namely VSV-G and S1 pseudotyped lentivirus particles. We observed comparable reduction in transduction when both pseudotyped lentivirus particles were treated with increasing amounts of LL37 (Fig S1E).

**Figure 1.**
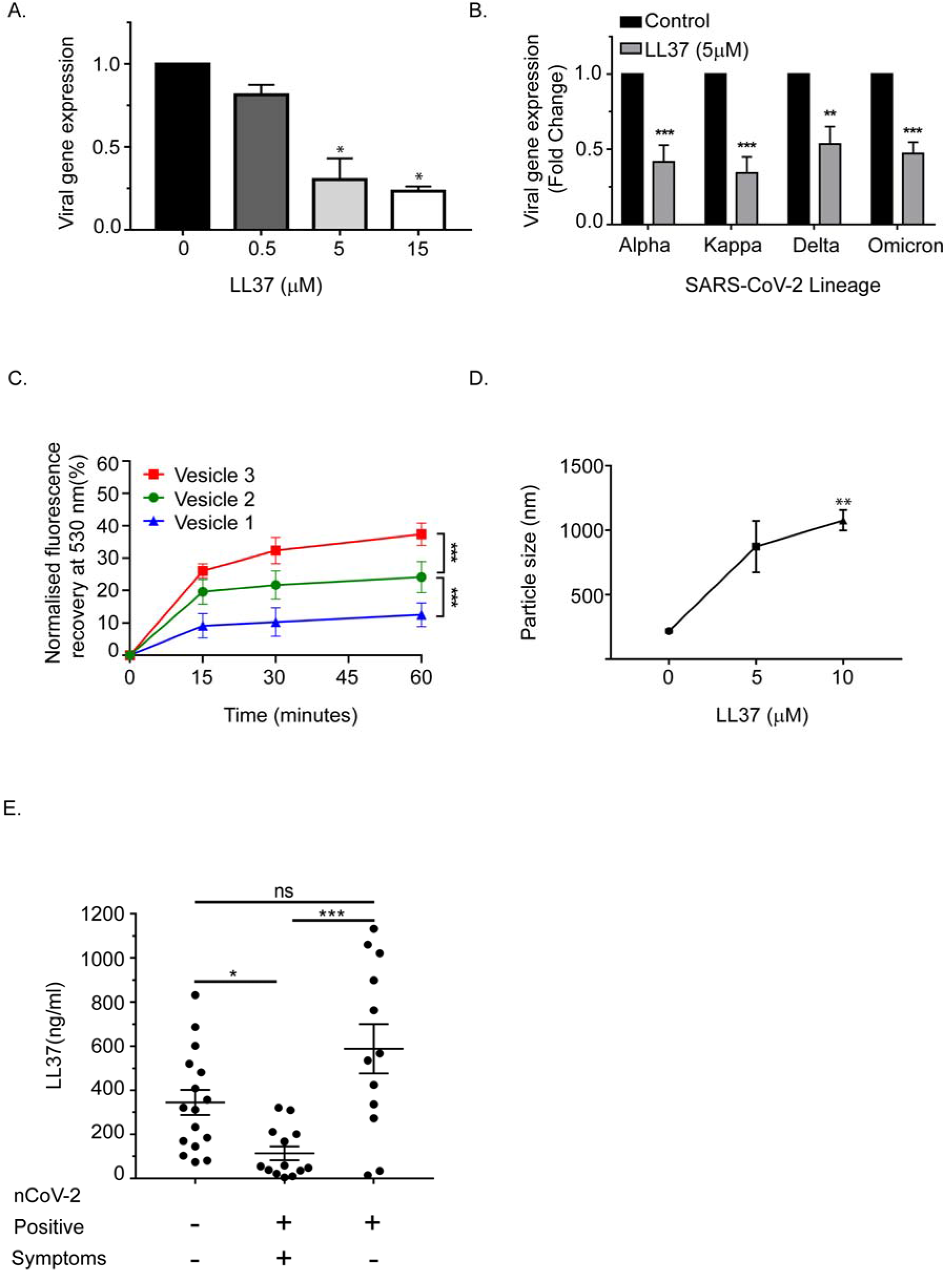
Antiviral activity of LL37 against SARS-CoV-2 variants. (A) Effect of increasing LL37 concentrations on SARS-CoV-2 neutralization (qRT-PCR) (n=3) (B) Effect of LL37 on the infectivity of various SARS-CoV-2 variants (qRT-PCR) (n=3) (C) Membrane disruption assay using virus like vesicles, mimicking viral membrane and PS concentration (FRET) (n=3) (D) Particle size analysis of SARS-CoV-2 in the presence of LL37 (DLS) (n=3) (E) Measurement of LL37 in various patient cohorts (ELISA) (A total of 41 individuals were subdivided into negative (16), positive symptomatic (13), and positive asymptomatic (12) cohorts) [Data are shown as mean⍰±⍰SEM, statistical analysis was done using one-way ANOVA (A,D), two-way ANOVA (B,C), and Kruskal Wallis test (non-parametric ANOVA) (E), *p≤0.05, **p≤0.001, ***p≤0.0001]

These results are consistent with reports that LL37, a cationic peptide, can execute its antimicrobial activity by attacking the negatively charged membrane of pathogens^13^. Enveloped viruses such as coronavirus that assemble virions by budding off from the endoplasmic reticulum membrane have a negatively charged membrane due to a higher content of phosphatidylserine (PS)^14,15^. To mimic the membrane composition of a generic coronavirus, we prepared three different vesicles in which PS composition was varied according to the published range of ER-derived virions^15^ (Fig S1F). We observed that increasing the percentage of PS resulted in an increase in the negative surface charge on the vesicles (Fig S1F), which was neutralized by the presence of LL37 (Fig S1G). These results suggest that the positively charged peptide can coat the outer leaflet of the bilayer by electrostatic interactions. To determine the consequence of the interaction of LL37 with the vesicles, we assayed whether membrane integrity was compromised. Using a fluorescence resonance energy transfer (FRET) based membrane disruption assay (schematically shown in Fig S1H), we observed a reduction in FRET (fluorescence recovery at 530nm) when vesicles were treated with LL37 (Fig. 1C). These results indicate that LL37 is more effective in interacting with and disrupting membranes with a higher negative charge (Fig S1F). Previous reports have also indicated that disruption of vesicle membranes by positively charged polymers leads to vesicle clumping^16^. We therefore investigated whether disruption of the pseudoviruses and SARS-CoV-2 by LL37 would result in their aggregation leading to increase in particle size as measured by dynamic light scattering (DLS). Consistent with the reported effect of cationic polymers on negatively charged membranes, we observed an increase in particle size of SARS-CoV-2 (Fig 1D) as well as pseudotyped virus (VSV-G and Spike) (Fig S1I) upon treatment with LL37.

Disease severity of several viral respiratory infections has been inversely correlated with LL37 levels^17^. Since it has been shown that salivary burden of SARS-CoV-2 correlates with disease severity in patients^18^, we compared the levels of secreted LL37 in the saliva of SARS-CoV-2 infected and uninfected individuals (patient information in (Fig S1J)). Symptomatic individuals had on average ~3-fold less LL37 than uninfected individuals (Fig 1E). Interestingly, asymptomatic positive patients had equivalent levels of LL37 as uninfected individuals. These results suggest that lower LL37 levels may potentially render individuals more susceptible to a symptomatic infection.

Since lower levels of LL37 are associated with the symptomatic COVID-19 patient group (Fig 1E), we speculated that increasing the level of LL37 or enhancing the activity of the existing LL37 might serve as a potential means of combating SARS-CoV-2 infection. One method of enhancing the activity of LL37 is to decrease its inherent self-aggregation^12^ and thereby increase its bioavailability. A common approach to prevent aggregation is through the use of a hydrotrope such as niacinamide (vitamin B3). It is a generally regarded as safe (GRAS) substance used to increase the solubility, and therefore the activity, of various drugs^19^. Indeed, we observed that LL37 supplemented with niacinamide exhibited an enhanced potency against infection by different variants of SARS-CoV-2 (Fig 2A, S2A). To understand the mechanism of LL37 interaction with lipid membranes in the presence of niacinamide, we used atomistic molecular dynamics (MD) simulations. Our simulations support the hydrotropic solubilisation of LL37 by an aqueous solution of niacinamide. We found that niacinamide transiently associated with mainly the hydrophobic and non-polar residues of LL37 (Fig S2B) by both aromatic-π and van der Waals interactions. These predominantly included phenylalanine (Phe5, Phe6, Phe17 and Phe27) and isoleucine (Ile20 and Ile24) residues (Fig 2B), which mediates LL-37’s self-aggregation and therefore reduced activity^20^. Our simulations suggest that encapsulating of aggregation prone residues of LL37 by niacinamide would likely improve the bioavailability of the peptide.

**Figure 2.**
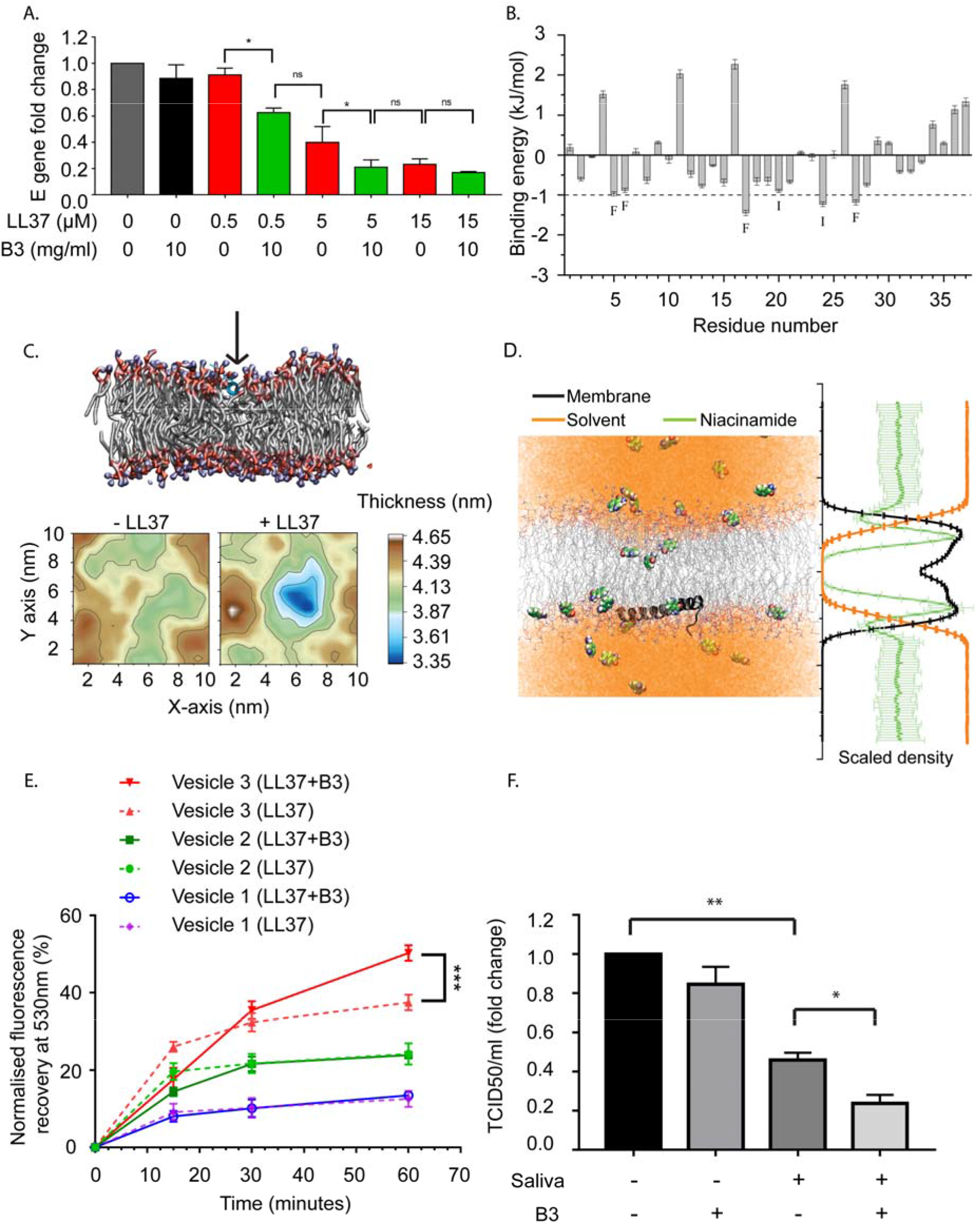
Effect of niacinamide on the antiviral activity of LL37. (A) Viral gene expression at different concentrations of LL37 in the presence of niacinamide (qRT-PCR) (n=3) (B) Contribution of each residue of LL37 towards interaction with niacianamide from MD simulations (mean of 3 independent simulations, SD propagated from each replicate). Residues contributing > 1 kJ/mol (*dotted line*) are labelled. (C) The LL37 peptide gets deeply embedded in the membrane in 200 ns (*top*). Representative membrane thickness averaged over the last 20 ns of MD simulations in absence and presence of the LL37 peptide (*bottom left and right respectively*). Color scale indicates membrane thickness in nm. (D) Representative snapshot (*left*) of niacinamide penetrating deeper than water into the hydrophobic core of the membrane (LL37, black peptide; lipid acyl chains, grey lines; niacinamide, space filling; water, orange) which is quantified on the *right*, averaging the molecular density (membrane, black; niacinamide, green; water, orange) over the entire 200 ns trajectory (5 replicates, mean ± SD) (E) Membrane disruption assay on addition of LL37 and niacinamide (FRET) (n=3) (F) Effect of niacinamide addition to saliva on SARS-CoV-2 neutralisation (TCID50) (n=4) [Data are shown as mean⍰±⍰SEM, Statistical analysis was done using student t-test for (A), two-way ANOVA for (E), and one-way ANOVA for (F), *p≤0.05, **p≤0.001, ***p≤0.0001]

In addition, we employed molecular dynamics simulation to investigate the early steps of LL37 adsorption on viral envelope-like membranes in the presence of niacinamide. These simulations demonstrate the lipid acyl chains and headgroups interacting with the LL37 peptide (Fig 2C, *top*). In accordance with previous literature^21^, these interactions pull the peptide deeper into the membrane resulting in local thinning in the bilayer and ultimately a dramatic destabilization of membrane (Fig 2C bottom, Fig S2C), with a concomitant destabilization of lipid ordering (Fig S2D). This configuration meant that the charged amino acid residues of LL37 faced the solvent and were free to interact with niacinamide (S2E). Additionally, this simulation predicts that niacinamide also penetrates into the membrane (Fig 2D) and may synergize with LL37 to disrupt the membrane. Thus, we conclude that niacinamide has dual activities: (i) hydrotropically increase the aqueous solubility of LL37, thereby rendering it more bioavailable and; (ii) cooperate with LL37 to destabilize the membrane.

To validate these computational results, we performed a FRET-based membrane disruption assay using artificial viral membranes (as described in Fig S1F, H) in the presence of LL37 and niacinamide. Consistent with simulation predictions, we observed that membrane disruption of liposomes by LL37 was enhanced in combination with niacinamide (Fig 2E), while niacinamide by itself was not antiviral. To test whether the effect of niacinamide can be reproduced with naturally produced AMPs, we analyzed its effect on AMPs that are highly secreted in saliva^22^ and from the skin^7^. We found that human saliva exhibits antiviral activity against SARS-CoV-2, which can be potentiated upon supplementation with niacinamide (Fig 2F). Likewise, we also observed that skin scrubs supplemented with niacinamide exhibited antiviral activity (Fig. S2C). Our body naturally synthesizes niacinamide, but interestingly, the biosynthetic pathways and precursor leading to niacinamide production are downregulated in symptomatic COVID-19 patients^23^. Thus, exogenous supplementation of niacinamide in symptomatic patients may potentiate the activity of naturally produced AMPs from the body’s epithelia.

The variants that exhibit increased transmissibility and disease severity are reported to contain mutations in the receptor binding domain (RBD) of the spike protein^24^. Given their key role in mediating viral entry into the host cell, many vaccines have been developed with antigens derived from the spike protein, and mutations pose a serious problem with vaccine escape^25^. In addition, the heavy glycan coating of the spike proteins can be a mechanism of camouflaging them from the immune system^26^. One approach to circumvent these problems is to target the viral envelope that originates from the host cell and is thus conserved among the different variants. Altogether, we show that the AMP LL37 has the potential to neutralise the SARS-CoV-2 viral infection by targeting its envelope and niacinamide further enhances this antiviral activity of the peptide. Our data on the symptomatic patient samples further substantiate this hypothesis and argues for an approach that would entail enhancing the efficacy of antimicrobial peptides for protection against viral infections. Therefore, either exogenous administration of the AMP with niacinamide or other strategies to boost the endogenous production of the peptide in combination with niacinamide could be a potent method to not only block viral transmission, but may be an effective therapy to limit viral load and disease severity of a patient post infection.

## Materials and Methods

### Cell lines and reagents

Caco2 (ATCC HTB-37), Vero-E6 (ATCC CRL-1586), Calu-3 (ATCC HTB-55) and HEK 293T cells (ATCC CRL-3216) were grown in Dulbecco’s Modified Eagle’s Medium (DMEM, Gibco), with 10% fetal bovine serum (FBS, Gibco), 100U/ml penicillin and 100μg/ml streptomycin (Gibco) at 37° C in 5% CO_2_ incubator. Human keratinocyte cell line (HaCaT cells) were grown and differentiated as described elsewhere^27^. All the experiments and incubations were carried out in serum free media such as OptiMEM (Gibco) or EpiLife (Gibco).

### Plasmids

Lentivirus packaging plasmid psPAX2 (Addgene #12260) and pMD2.G (Addgene #12259) were used for VSV-G pseudovirus production. For generating SARS-CoV-2 spike pseudotyped lentivirus, pTwist-EF1alpha-SARS-CoV-2-S-2xStrep (Addgene #141382)^28^ was used. pZip-mEF1a-ZsGreen-Puro was used for the ZsGreen expression.

### Transfection and pseudotype virus production

293T cells were transfected with psPAX2, pZIP-mEF1a-ZSGreen-Puro, and either pMD2.G or pTwist-EF1alpha-SARS-CoV-2-S-2xStrep using LTX (Invitrogen) transfection reagents according to the manufacturer’s protocol. Following a 48 hr transfection, the virus particle-containing media was harvested and virus particles were purified using Polyethylene glycol 8000 (Sigma).

### Transduction and FACS

About 0.1 to 0.2 MOI (Multiplicity of Infection) equivalent of VSV-G and pseudotype SARS-CoV-2 spike virus particles were treated with recombinant LL37 (Tocris) and niacinamide (vitamin B3) (provided by Unilever), both solubilised in sterile distilled water, for three hours at 37°C (in 100 uL volume). Treated lentivirus were then added to Caco2 cells (96 well plate with about 30,000 cells/well) for transduction. ZsGreen expression was monitored after 72 hr of transduction. The cells were trypsinized and single cell suspension was analysed in an Attune NxT flow cytometer with appropriate gates to determine percentage of transduction in each sample.

### Dynamic light scattering analysis

Particle Size Analyser: Litesizer 500 model with a scattering angle of 90° angle was used to measure the particle size of pseudovirus particles (VSV-G and SARS-CoV-2 spike). The particles were treated with LL37, Vitamin B3 or their combination for one hour at 37°C in 100 uL volume. The samples were then diluted to 1 ml with Epilife media (Cat No. MEPI500CA). Readings were taken in a quartz cuvette at 25°C in series for triplicates. The measured data were processed by Kalliope™ software and intensity distribution curve was generated with peaks on the y-axis corresponding to particle intensity and their respective size (in nm) represented on the x-axis.

### Preparation and biophysical characterization of liposomes

Vesicle 1 was prepared with molar ratio of 85% DPPC (Dipalmitoyl phosphatidylcholine), 15% DOPE (Dioleoyl phosphatidylcholine), 0% DOPS (1,2-dioleoyl-sn-glycero-3-phospho-L-serine), Vesicle 2 (DPPC: DOPE: DOPS: 83%: 11%: 6%) and Vesicle 3 (DPPC: DOPE: DOPS: 79%: 11%: 10%), at a total concentration of 0.5mM. Dry thin films of lipids were prepared in glass vials, followed by overnight hydration with deionized water to produce hydrated films. Vortexing vials for 2-3 min produced multilamellar vesicles. Subsequently, bath sonication (Qsonica sonicator, Q700) produced small unilamellar vesicles.

### Zeta Potential measurement

The vesicles’ sizes and zeta potentials (surface charges) were measured by photon correlation spectroscopy and electrophoretic mobility on a Litesizer 500 (Anton Paar). The sizes and potentials of vesicles were measured in deionized water with a sample refractive index of 1.32 and a viscosity of 0.6912 cP. The diameters of liposomes were calculated by using the automatic mode. The zeta potential was measured using the following parameters: dielectric constant 74.19, approximation (Smoluchowski) 1.50; maximum voltage of the current 100 V. All the measurements were taken in triplicates.

### FRET assay for membrane disruption

Vesicles 1, 2, and 3 (of concentration 0.5mM) were prepared with FRET pair Lipids NBD-PE and N-Rho-PE (Avanti-Polar Lipids, USA) were used as the donor and acceptor fluorescent lipids, respectively, with 0.005 mM NBD-PE and N-Rho-PE lipids (i.e., 1% with respect to the vesicle formulation content). Labelled vesicle formulations were placed in a fluorimeter, Microplate Fluorescence Reader (Horiba Instruments, USA) at 25° C and LL37 (5μM) was added in the one set of experiments. In another set of experiments, both Vitamin B3 (10mg/ml) and LL37 (5μM) were added to labelled vesicles., Fluorescence intensities were recorded as a function of time with excitation at 485 nm and emission at 530 nm. Data were normalized by considering a 100% value, which was obtained after lysing the liposomes with 1% TritonX-100. It was determined from the NBD-PE fluorescence intensity of the labelled vesicles formulation (experimental schematic in Fig S1H).

### SARS-CoV-2 infection

Experiments were carried out with 2019-nCoV/Italy-INMI1 strain unless otherwise stated to have used alpha, kappa, delta, and omicron strains of SARS-CoV-2. All the virus-related experiments were carried out in the BLiSc biosafety level 3 (BSL-3) laboratory. Virus particles, equivalent to 0.1 MOI, were incubated with recombinant LL37 and Vitamin B3 or buffer control for three hours in 100 μl volume. The pre-treated particles were then added to the host cells in a 24 well plate with about 240,000 cells/well and allowed to adsorb for one hour. The cells were then washed and replenished with fresh media.

### RNA isolation

24 hr after incubation of pre-treated virus particles on cells, RNA was isolated using RNAiso Plus (Takara). 1 μg of RNA was used to prepare cDNA using the PrimeScript kit (Takara). cDNA equivalent to 100 ng of RNA was used for setting up the qPCR reaction using SYBR green master mix (Applied Biosystems). PCR reactions were performed using the CFX384 Touch Real-time PCR detection system (BioRad). Viral gene expression changes were calculated following normalization to β-actin using the comparative Ct (cycle threshold) method.

### ELISA

Conditioned media of Calu3 and differentiated HaCaT were collected from 72 hr old confluent culture plates. The media were centrifuged at 10,000 × g for 3 minutes and used for ELISA. Unstimulated saliva samples were collected from uninfected donors. The samples were cleared by centrifugation at 10,000 × g for 5 minutes and the concentration of LL37 was determined by ELISA using a commercially available analysis kit specific for LL37 (Hycult Biotech, HK321-02), following manufacturer’s protocol. The absorbance was measured at 450 nm using a microplate reader (Varioskan, Thermo).

### TCID50

A median tissue culture infectious dose (TCID50) assay was performed to identify the viral titer of SARS-CoV-2 in control and treated conditions. VeroE6 cells were cultured in a 96-well tissue culture plate and varying dilutions of the virus were added. The virus was allowed to adsorb for 1 hr, following which cells were thoroughly washed and media was replenished. After incubation for 72 hr, the percentage of infected wells was observed for each dilution. These results were used to calculate the TCID50 value using a TCID50 calculator (by Marco Binder; adapted at TWC).

### Atomistic molecular dynamics simulations

Unrestrained all atom molecular dynamics (MD) simulations were performed on two different systems. The first system comprised one copy of LL37 (PDB id: 2K6O), 40 molecules of niacinamide (3NAP from CGenFF^29^) in a water solvated box of 1000 nm^3^ with 150 mM NaCl (Figure 2B, S2B) which corresponds to 1 LL37 peptide molecule in a 10 mg mL^−1^ nicotinamide solution. This system was simulated in triplicates of 50ns each.

The other simulation was composed of a symmetric coronavirus lipid bilayer^15^ comprising POPC:POPE:POPS:cholesterol in a 70:15:10:5 ratio. One copy of LL37 peptide (2K6O) was adsorbed on the surface of the membrane using the initial configuration from the PPM server^30^ and 40 molecules of niacinamide (3NAP, CGenFF) were added in the box of 10 × 10 × 14.5 nm^3^ (Figure 2C, S2C-E). This was solvated and sodium counter ions were added to neutralize the charge. This configuration of 138,332 atoms was simulated in 5 replicates for 200 ns each. As a control, a fully solvated identical membrane was simulated in duplicate for 100ns each. All the particles were described by the CHARMM36 force field^31^ and the isothermal-isobaric production simulations were performed using Gromacs^32^ 2018. Inbuilt gromacs routines and g_mmpbsa^33^ were used to analyze the simulations. Area per lipid and membrane thickness were estimated using Gridmat^34^ and the lipid order (*S*_*CD*_) was determined from equation 1 as implemented in Membrainy^35^.

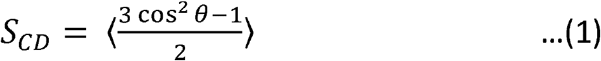

### Saliva and skin scrub treatment of SARS-CoV-2

Virus particles equivalent to 0.1MOI were incubated with 100μl of either skin-scrub (5x concentrated with 3kDa Amicon filters) or control buffer at 37°C for four hours. Following this, the pre-treated particles were added on VeroE6 cells for TCID50 assay, or Caco2 cells for qPCR and were allowed to adsorb for one hour at 37°C. The cells were then washed and replenished with fresh media.

## ACKNOWLEDGEMENTS

The authors would like to thank Jamora lab members for their critical review of the work and insightful discussions. We would like to thank Shah-E-Jahan (VS lab) for the help with SARS-CoV-2 viral culture, Johan Ajnabi (CJ lab) and Shreyaa S. (CJ lab) for the help with cell culture maintenance and pseudovirus generation. We also acknowledge BLiSc biosafety level 2 and 3 facilities, and high performance computing cluster at NCBS.

This work was supported by a grant from the Department of Biotechnology of the Government of India, Institute for Stem Cell Science and Regenerative Medicine (inStem) core funds, and financial support from Unilever. S.U.K and B.D are supported by core funds at inStem.

---

**Figure S1.**
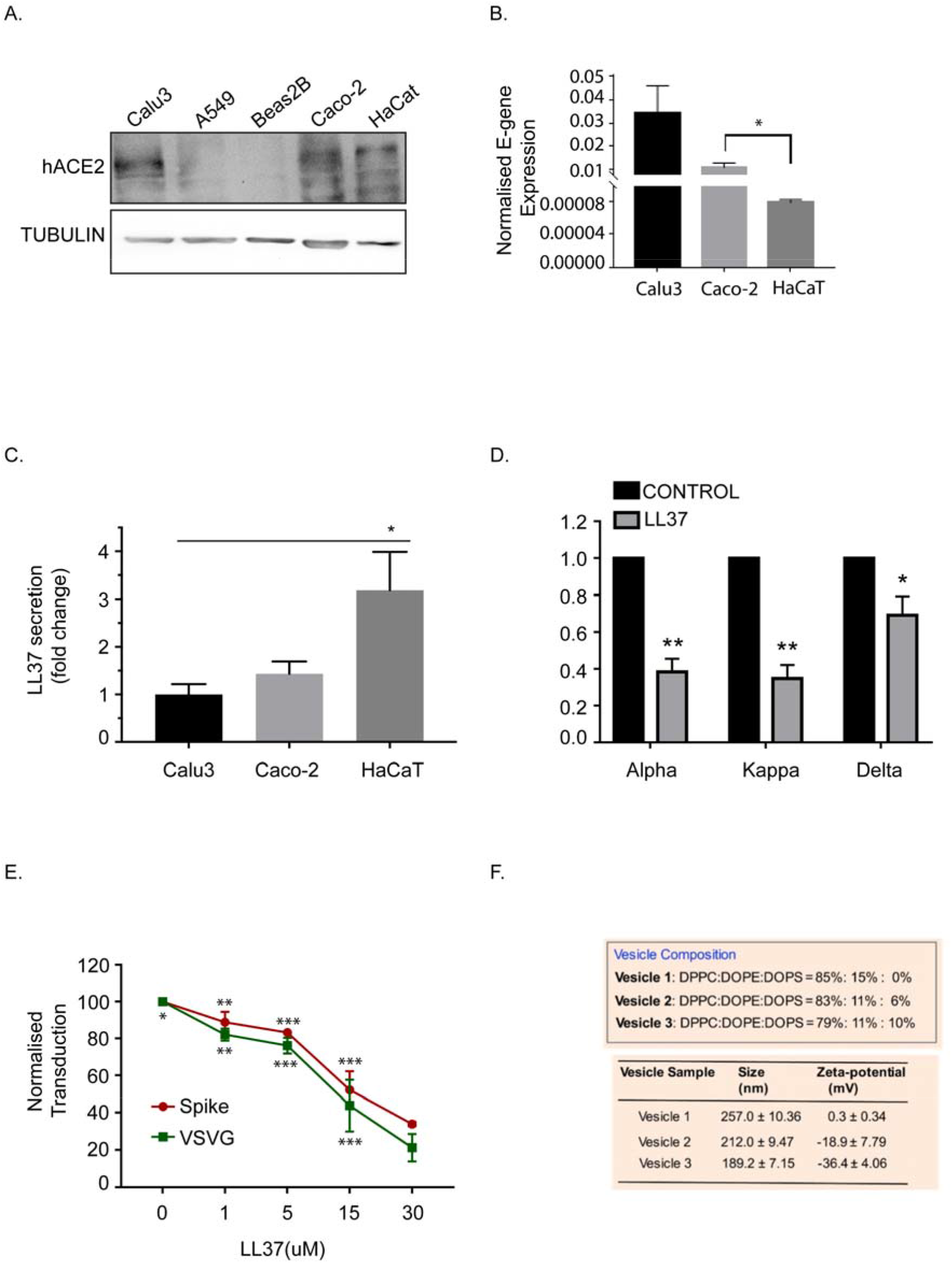
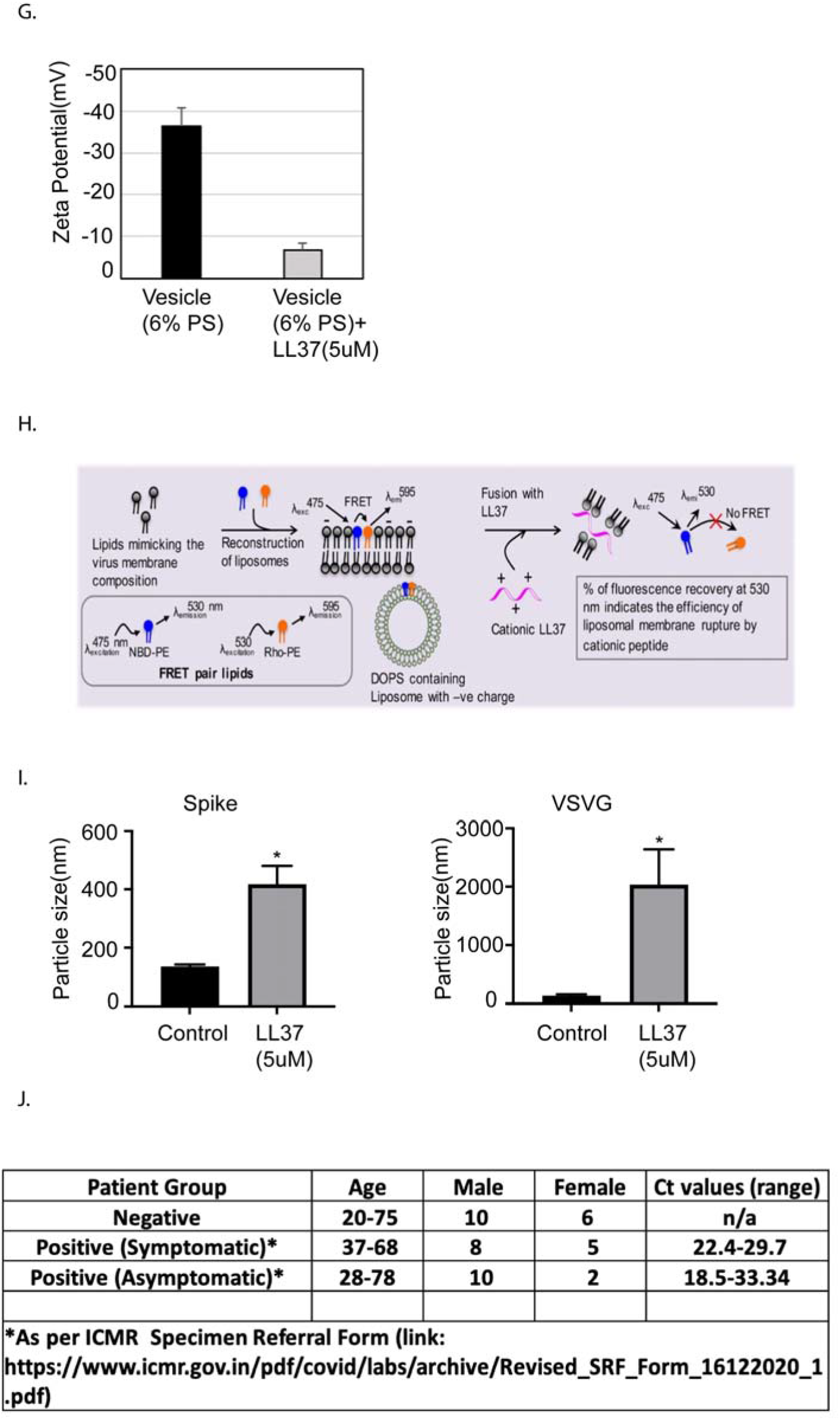
(A) Ace2 protein levels in various epithelial cell lines (Western blotting) (n=2) (B) SARS-CoV-2 E-gene expression in Calu3, Caco2, and HaCaT cells, 24 hr post infection (qRT-PCR) (n=2) (C) Secreted LL37 levels in Calu3, Caco2 and HaCaT cells (ELISA) (n=4) (D) Effect of LL37 on the TCID50/ml of various SARS-CoV-2 strains (n=3) (E) Neutralization of Pseudo type virus (spike and VSVG) in the presence of LL37 (FACS) (n=3) (F) Membrane composition, size, and charge of virus like vesicles (G) Charge neutralization of vesicles in the presence of LL37 (n=3) (H) Principle and methodology of FRET based membrane disruption assay (I) Particle size analysis of Pseudovirus (Spike and VSVG) in the presence of LL37 (DLS) (n=3) (J) Details of the patient group considered for this study [Statistical analysis was done using Student’s t-test (B,G, I), one-way ANOVA (C), and two-way ANOVA (D,E), *p≤0.05, **p≤0.001, ***p≤0.0001]

**Figure S2.**
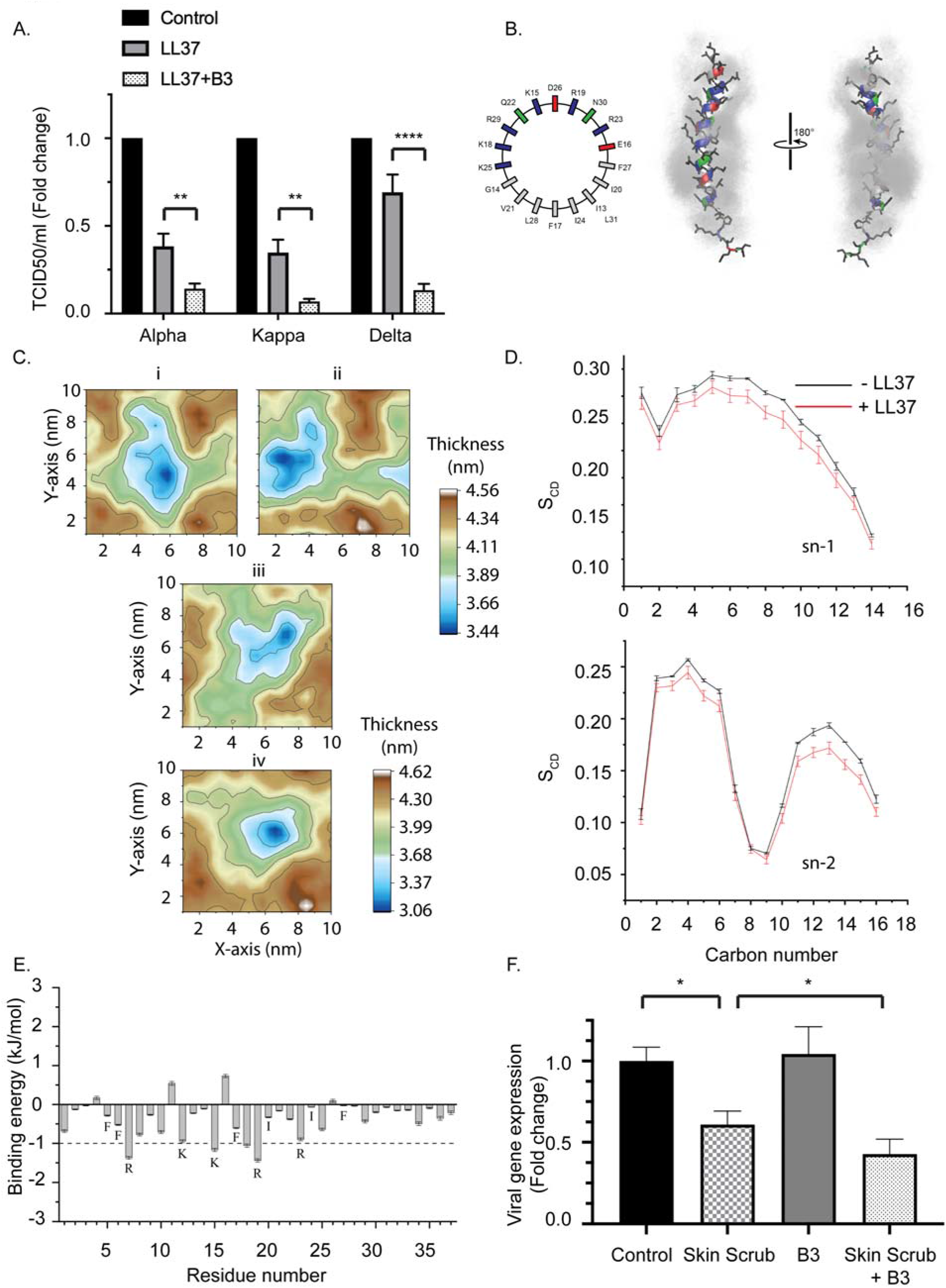
(A) Effect of niacinamide combined with LL37 on the TCID50/ml of various SARS-CoV-2 strains (n=3) (B) Partial sequence of the amphipathic LL37 peptide is represented as an idealized helical wheel (left) with residues colored according to their nature (hydrophobic, grey; polar, green; acidic, red; basic, blue). Two views of the LL37 peptide (color scheme same as the wheel; sidechains, black) overlaid with niacinamide molecules (grey) within 6 Å of the peptide (over 10 ns). The 10 ns overlay emphasizes the solvent-like encapsulation of the peptide by niacinamide with greater residence over the hydrophobic face of the helix than the polar/charged face (C) Mean thickness of the membrane averaged over the last 20 ns of four simulations (i-iv). The adsorbed LL37 peptide caused a deformation of > 1 nm on the membrane (i-iii) and > 1.5 nm in one case (iv). Color scale indicates membrane thickness in nm (D) The LL37 peptide gets deeply embedded in the membrane in 200 ns (Fig 2C (*top*)) which causes a reduction in lipid ordering. Both the saturated (sn-1) and the unsaturated (sn-2) lipid acyl chains exhibit reduced ordering in presence of the LL37 peptide, quantified using the lipid order parameter (S_CD_) (mean ± SD from last 20ns of 5 independent simulations) (E) Niacinamide in the membrane forms hydrogen bonds with the Lys (K) and Arg (R) residues of LL37. Whereas, the aliphatic residues of LL37 which preferentially interacted with niacinamide in solution (Fig 2B) are now predominantly engaged with the hydrophobic acyl chains and so, are unable to contribute much to bind niacinamide (F) Effect of skin scrub and its combination with niacinamide on SARS-CoV-2 viral gene expression (qRT-PCR) (n=3) [Statistical analysis was done using two-way ANOVA for (A) and student t-test for (F). *p≤0.05, **p≤0.001, ***p≤0.0001]

